# Intrinsically disordered proteins and structured proteins with intrinsically disordered regions have different functional roles in the cell

**DOI:** 10.1101/646901

**Authors:** Antonio Deiana, Sergio Forcelloni, Alessandro Porrello, Andrea Giansanti

## Abstract

Many studies about classification and the functional annotation of intrinsically disordered proteins (IDPs) are based on either the occurrence of long disordered regions or the fraction of disordered residues in the sequence. Taking into account both criteria we separate the human proteome, taken as a case study, into three variants of proteins: i) *ordered proteins* (ORDPs), ii) *structured proteins with intrinsically disordered regions* (IDPRs), and iii) *intrinsically disordered proteins* (IDPs). The focus of this work is on the different functional roles of IDPs and IDPRs, which up until now have been generally considered as a whole. Previous studies assigned a large set of functional roles to the general category of IDPs. We show here that IDPs and IDPRs have non-overlapping functional spectra, play different roles in human diseases, and deserve to be treated as distinct categories of proteins. IDPs enrich only a few classes, functions, and processes: *nucleic acid binding proteins, chromatin binding proteins, transcription factors*, and *developmental processes.* In contrast, IDPRs are spread over several functional protein classes and GO annotations which they partly share with ORDPs. As regards to diseases, we observe that IDPs enrich only cancer-related proteins, at variance with previous results reporting that IDPs are widespread also in cardiovascular and neurodegenerative pathologies. Overall, the operational separation of IDPRs from IDPs is relevant towards correct estimates of the occurrence of intrinsically disordered proteins in genome-wide studies and in the understanding of the functional spectra associated to different flavors of protein disorder.

## Introduction

Over the last two decades, the concept of *‘intrinsic disorder’* has emerged as a prominent and influential topic in protein science [1-6]. The discovery of intrinsically disordered proteins (IDPs), i.e., functional proteins lacking a well-defined tertiary structure, has challenged the traditional sequence-structure-function paradigm [7,8]. IDPs constitute, in eukaryotes, a substantial part of the cellular proteome and are involved in many biological processes that complement the functional repertoires of ordered proteins [9,10]. Ever-increasing experimental evidence has revealed the presence of disordered regions also in well-structured proteins [11,12]. Then the distinction between intrinsically disordered proteins, i.e., proteins lacking a tertiary structure, and structured proteins containing intrinsically disordered regions gradually emerged [13-16]. An operational distinction of these two variants is the main objective of the present work. A note on terminology is in order at this point. The editors of the journal Intrinsically Disordered Proteins made an effort to disambiguate the semantics of protein intrinsic disorder [17]. They proposed “intrinsically disordered proteins” as a unifying term, recognizing that it is a compromise “far from being ideal”, and suggesting that “additional descriptors” would emerge, apt to clarify the many aspects of “structurelessness” that are included in one term. The terminology we adopt here is surely also not ideal. Let us try to make clear that we use here IDPs, when referring to previous studies, as the general unifying term proposed by Dunker et al. We use IDPs also to denote a subgroup of Dunker et al.’s IDPs, a variant that is operationally specified here as distinct from the variant of mostly structured proteins with long intrinsically disordered regions (we call here IDPRs).

The biological role of IDPs has been the focus of a growing number of publications, in the recent past. Xie et al., for example, in a series of three articles compiled an anthology of the functional roles of IDPs [18-20]. It has been reported that out of the 710 keywords recording biological functions in the Uniprot/Swissprot database, 310 (44%) are associated with ordered proteins and 238 (34%) with IDPs [21]. Several studies have reported lists of Gene Ontology functional classes that are enriched in IDPs, including but not limited to: cell regulation, transcription, translation, signaling, and alternative splicing [22-29]. Different studies highlight the occurrence of unfolded regions in proteins that function as chaperones for other proteins or RNA molecules, transcription factors, effectors, assemblers, and scavengers [13-15,26]. Importantly, many proteins with disordered regions have been associated with human diseases [8,30,31], and a D2 concept (disorder in disorders) emerged [32]. However, in all these studies what we distinguish here as IDPs and IDPRs were considered as a unique variant, possibly aggregating proteins which are structurally and functionally different. This is still reflected, for example, in a recent paper by Darling and Uversky [33]. IDPRs, as we define them, may have a well-defined 3D structure that either can undergo conformational, allosteric changes (e.g., after a post-translational modification [34]) or accommodate functional, locally disordered domains, that are limited to less than 30% of the structure. The presence of unstructured loops or domains, e.g., linear motifs and Molecular Recognition Features (MORFs), can be important for these proteins to target low-affinity substrates enlarging the repertoire of protein interactions [14].

Let us remind that one of the features that distinguishes IDPs from structured proteins is their different interaction mechanism with target substrates. According to the classic *lock-and-key mechanism* [35], the specificity and affinity required for molecular interactions depend on the complementarity of the binding interfaces. However, even proteins that are disordered under physiological conditions or contain long unstructured regions can form stable complexes [8,36]. In this context, the lack of a well-defined 3D-structure represents a major functional advantage for IDPs, allowing them to interact with a broad range of substrates with relatively high-specificity and low-affinity, often undergoing a disorder-to-order transition upon binding [2,8,26,37,38]. However, despite that this description makes sense for proteins lacking a tertiary structure (IDPs), it is reasonable to suppose that structured proteins with long disordered regions (IDPRs) still interact with substrates through a mechanism similar to the lock-and-key. It has been reported that disordered regions have many post-translational modification sites (PTMs) [8,29,39-41], which induce conformational changes leading to *one-lock-many-keys interactions* [33]. A recent study reviews different interaction mechanisms for unfolded and folded proteins [34]. Unfolded proteins are characterized by *induced-fit interactions*, whereas *conformational change interactions* characterize folded proteins. Although the recent literature does not explicitly separate IDPs from IDPRs, it is reasonable to hypothesize that IDPRs typically undergo conformational changes after post-translational modifications, whereas high-specificity and low-affinity induced fit mainly characterize IDPs.

Many studies aiming at a general classification of IDPs’ functional roles were based on the required presence of long disordered regions in the sequence [18-20, 42-45]. However, the presence of a long disordered segment does not necessarily imply that a protein lacks a three-dimensional structure. In support of this claim, Gsponer et al. showed that also the percentage of disordered residues plays an important role in determining the half-life of a protein [46]. Therefore, both the length of disordered domains and the overall fraction of disordered residues in the sequence are critical parameters to distinguish different types of protein disorder.

Following clear-cut criteria inspired by experimental observations [14,46], we compiled two lists of human IDPs and IDPRs. We identified IDPs as those proteins having more than 30% of disordered residues (implying a shorter half-life [46]). We defined IDPRs as proteins with at least one long disordered segment (suggesting a higher sensitivity to proteolytic degradation [14]) but less than 30% of disordered residues in their sequence. We have assessed both the over-representation and the enrichment of IDPRs and IDPs in different functional classes and gene ontologies. Moreover, we have investigated the relative occurrence of IDPRs and IDPs in proteins related to cancer, cardiovascular, and neurodegenerative diseases.

Our results indicate that IDPRs and IDPs are structurally different, have different functional spectra, and play different roles in human diseases.

## Materials and Methods

### Dataset of protein sequences

We downloaded the human proteome from the UniProt-SwissProt database (http://www.uniprot.org/uniprot) [47], release 2018_07. We selected 20386 human proteins by searching for reviewed proteins belonging to Homo Sapiens (Organism ID: 9606, Proteome ID: UP000005640).

### Dataset of protein sequences associated to diseases

To select the proteins associated to cancer, cardiovascular, and neurodegenerative diseases, we used the lists of keywords found in Iakoucheva *et al.* [48], Cheng *et al.* [49], and Uversky [50]. Then, we extracted the proteins annotated with those keywords from the UniProt-SwissProt database. The lists of disease-related proteins are available as Supplementary Materials.

### Prediction of protein disorder

We identified disordered residues in the protein sequences using MobiDB (http://mobidb.bio.unipd.it) [51], a consensus database that combines experimental data (especially from X-ray crystallography, nuclear magnetic resonance (NMR) and cryo-electron microscopy (Cryo-EM)), curated data and disorder predictions based on various methods. From the initial dataset, we discarded proteins not annotated in MobiDB. Of the 20386 initial proteins, we analyzed 20030 proteins, i.e. 98.3% of the initial dataset.

### Identification of IDPs and IDPRs

To partition the human proteome into different protein variants, we used two parameters: the percentage of disordered residues and the length of the longest disordered domain in the sequence. Based on these two parameters, we defined:

i. *ordered proteins* (ORDPs): they have less than 30% of disordered residues, no C- or N-terminal segments longer than 30 consecutive disordered residues as well as no segments longer than 40 consecutive disordered residues in positions distinct from the N-and C-terminus;
ii. *proteins with intrinsically disordered regions* (IDPRs): they have less than 30% of disordered residues in the polypeptide chain and at least either one C- or N-terminal segment longer than 30 consecutive disordered residues or one segment longer than 40 consecutive disordered residues in positions distinct from the N- and C-terminus;
iii. *intrinsically disordered proteins* (IDPs): they have more than 30% of disordered residues in the polypeptide chain.

Generally speaking, ORDPs are proteins with a limited number of disordered residues and the absence of disordered domains. IDPRs, unlike ORDPs, are proteins with at least one long disordered segment (implying a short half-life [14]) accommodated in globally folded structures. The rationale of taking into account the location of the long disordered segments in the definitions i) and ii) is discussed below in the discussion section. IDPs are proteins with a significant percentage of disorder (implying an unfolded structure and a high susceptibility to proteolytic degradation [46]).

### Clustering of the variants of disorder through functional profiles

The functional profiles of ORDPs, IDPRs and IDPs were defined considering several annotations, such as: biological processes, cellular components and molecular functions from the Gene Ontology database [52] and protein classes from the PANTHER database [53-55].

Functional profiles associated with each protein variant are unitary vectors whose components are estimated as the number of proteins of a variant in each functional class divided by the number of proteins in that specific variant. Functional distances between protein variants were evaluated through the Euclidean distance of their profiles. The protein variants were then clustered using the average linkage hierarchical clustering algorithm [56]. The stability of the clustering against noise in the attribution of proteins to the different functional classes was assessed through bootstrap [57]. In each bootstrap re-sampling, new lists of ORDPs, IDPRs, and IDPs were compiled by randomly picking-up (with repetition) proteins in each variant. The robustness of two variants clustering together was estimated by *f*, the number of times this happened, divided by the number of bootstrap replicates [58]. Similarly, we evaluated the probability that two variants cluster just by chance by evaluating the fraction *f* of times that the two variants are clustered using resampled lists of ORDPs, IDPRs, and IDPs in which proteins from the entire human proteome are randomly inserted.

### Over-representation of a protein variant in a functional class

The fractions of ORDPs, IDPRs, and IDPs were evaluated in each functional class. A protein variant is over-represented in a functional class if the frequency of its proteins in such a class is higher than the frequency of the same variant in the human proteome. The statistical significance of the differences was assessed through a binomial test [53, 59]. We did three statistical analyses, one for each protein variant. A conservative p-value, accounting for the multiple testing, was obtained through Bonferroni correction [60].

### Enrichment of a protein variant in a functional class

A functional class is enriched in the protein variant that has the highest frequency and depleted in the protein variant that has the lowest frequency. We evaluated the statistical significance of enrichment/depletion through a cascade of tests. Firstly, we tested the null-hypothesis that the three variants (ORDPs, IDPRs, and IDPs) have the same frequencies in a functional class. Through a goodness-of-fit (chi-square) test we rejected the null hypothesis when the p-value is lower than 0.05. Secondly, we tested the two pairs of variants with the highest and the two pairs with the lowest frequencies with two further goodness-of-fit tests of the hypothesis of equal probabilities. In each of the two tests, if the p-value is lower than 0.05 then the hypothesis of equal frequency of the pairs is discarded. Eventually we concluded that a given functional class is enriched in the protein variant that has the highest significant frequency and depleted in the protein variant that has the lowest significant frequency. Looking at Table in S6 Table it is clear that, if the first test is not passed no conclusion on the enrichment can be made. Once the first test is passed then three cases are possible in each functional class: i) neither significant enrichment nor depletion; ii) either significant enrichment or significant depletion; iii) both significant enrichment and significant depletion.

## Results

### ORDPs, IDPRs and IDPs in the human proteome

In Table 1 we report numbers and frequencies of ORDPs, IDRPs, and IDPs in the human proteome. Most human proteins are ORDPs, followed by IDPs and IDPRs. The frequency of IDPs (proteins with more than 30% of disordered residues) is about 32%, a result consistent with Colak *et al.* [61]. According to previous estimates [62], the percentage of all proteins containing disorder (IDPRs and IDPs) is roughly 51%. Therefore, the classification of IDPRs either as ordered or disordered is important to correctly estimate the overall percentage of protein intrinsic disorder in the human proteome.

**Table 1.** Number and frequency of ORDPs, IDPRs and IDPs in the human proteome.

| Variant | Number of proteins | % |
| --- | --- | --- |
| ORDPs | 9702 | 49 |
| IDPRs | 3862 | 19 |
| IDPs | 6466 | 32 |
| **Total** | **20030** | **100** |

### Protein variants: distance of functional profiles

Functional distances between protein variants are reported in Fig 1 as hierarchical trees (distance matrices are reported in Table in S1 Table) relative to gene ontology biological processes, cellular components, molecular functions, and to the PANTHER protein classes. IDPRs robustly cluster together with ORDPs. We conclude that, as a general trait, IDPRs have functional profiles which are more similar to those of ORDPs than to those of IDPs.

**Fig 1.**
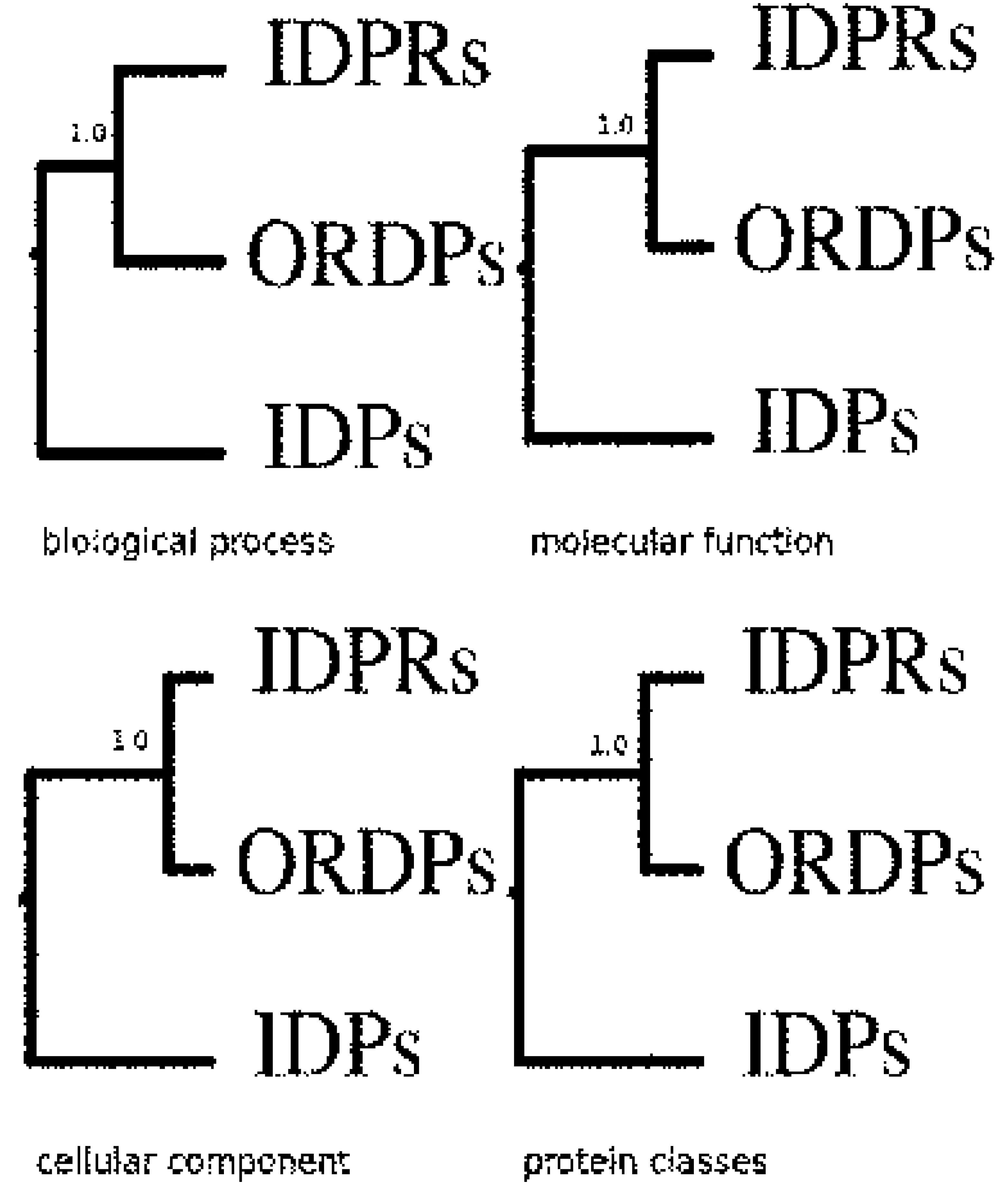
Clusters of the protein variants in the human proteome. Hierarchical clusters of ORDPs, IDPRs, and IDPs, based on different functional profiles. ORDPs and IDPRs are robustly separated, as indicated by bootstrap scores, the number of times these two groups clustered together divided by the number of bootstrap replicates (1000).

To further check that ORDPs and IDPRs do not cluster together by chance, we compared the hierarchical clusters of ORDPs, IDPRs, and IDPs in Fig 1 with the clusters obtained by repeatedly resampling each variant with proteins randomly selected from the human proteome. *f* values are comprised between 0.25 and 0.37 (see Figure in S1 Text). We, therefore, concluded that the lower distance between the functional profiles of ORDPs and IDPRs, observed in the human proteome (Fig 1), is due to functional specialization and not by chance.

### Protein variants: over-representation in the functional classes

A protein variant is over-represented (under-represented) in a functional class if its frequency is significantly higher (lower) than the frequency it has in the human proteome. In Fig 2 and in Table in S2 Table, we report the over- (under-) representation of ORDPs, IDPRs, and IDPs in the functional classes of PANTHER.

**Fig 2.**
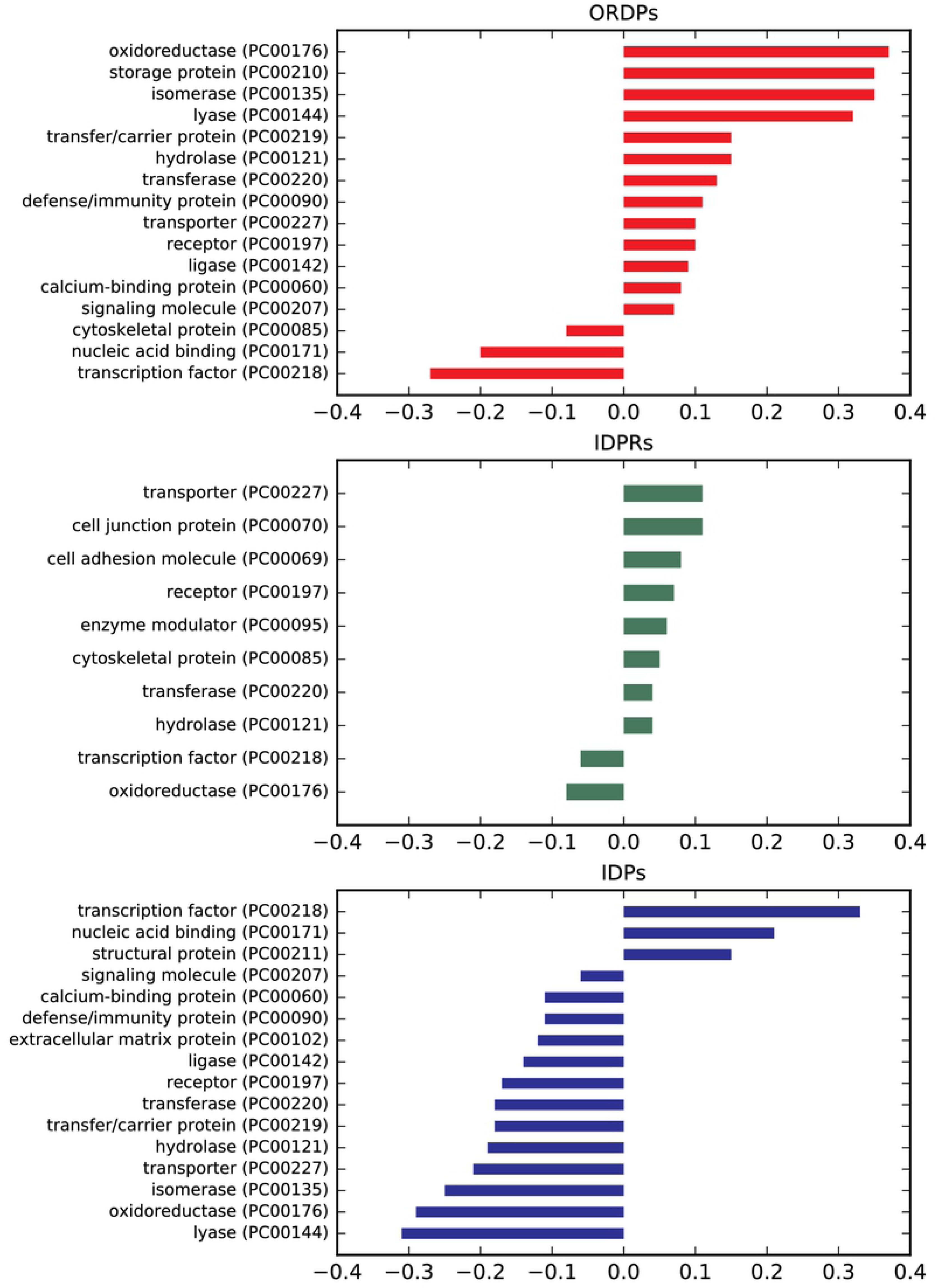
Over-(under-) representation of the protein variants in the PANTHER protein classes. Bar charts of the normalized differential occurrence of ORDPs, IDPRs, and IDPs in various protein classes, with respect to the human proteome (the reference). Only statistically significant differences are reported (p-value <0.05).

As expected, ORDPs are over-represented among *enzymes* (oxidoreductases, isomerases, lyases, hydrolases, transferases, ligases), and in several other protein classes related to *storage, defense/immunity, transporters, receptors, transfer/carriers and calcium-binding proteins*. ORDPs are under-represented among cytoskeletal proteins, nucleic acid binding proteins and transcription factors.

IDPRs are over-represented among *transporters, cell-junction, cell adhesion, receptors, enzyme modulators, cytoskeletal proteins*. Moreover, they are over-represented in *enzymes* such as hydrolases and transferases. IDPRs are under-represented among transcription factors and oxidoreductases. Notably, both ORDPs and IDPRs are over-represented in transporters, receptors, hydrolases, and transferases; interestingly, IDPs are under-represented in all these latter functional classes.

IDPs are over-represented only among *structural proteins, transcription factors*, and *nucleic-acid binding proteins*; note that the two latter functions are not over-represented both in ORDPs and IDPRs.

We also compared the over- (under-) representation of the variants in Gene Ontology annotations related to biological processes, molecular functions and cellular components (for the details see Figures in S1, S2, S3 Figures and Tables in S3, S4, S5 Tables).

In a nutshell, it is worth noting that each variant displays articulated spectra of specific GO annotations, that are generally consistent with the closeness of IDPRs to ORDPs already shown above, based on functional distances and PANTHER classes. On the contrary, IDPs have a well characterized, peculiar, set of annotations. Indeed, IDPs are specifically over-represented in molecular functions related to *nucleic acid binding* and *chromatin binding*, and *developmental biological processes* (i.e., pattern specification processes, ectoderm, embryo, mesoderm and system development), where ORDPs are under-represented and IDPRs are not significantly represented.

### Enrichment of functional classes in protein variants

In each protein functional class, the three protein variants (ORDPs, IDPRs, IDPs) are differently distributed (Fig 3). A functional class is enriched (depleted) in the variant that has the highest (lowest) relative frequency. The statistical significance of the enrichment (depletion) is evaluated with a specific test (see Materials and Methods). In Fig 3 and Table in S6 Table, we report the protein classes that reached statistical significance in the assessment of their enrichment (depletion). Only two protein classes are enriched in IDPs: *nucleic acid binding proteins* and *transcription factors*. The other classes are enriched in ORDPs. No protein class is enriched in IDPRs.

**Fig 3.**
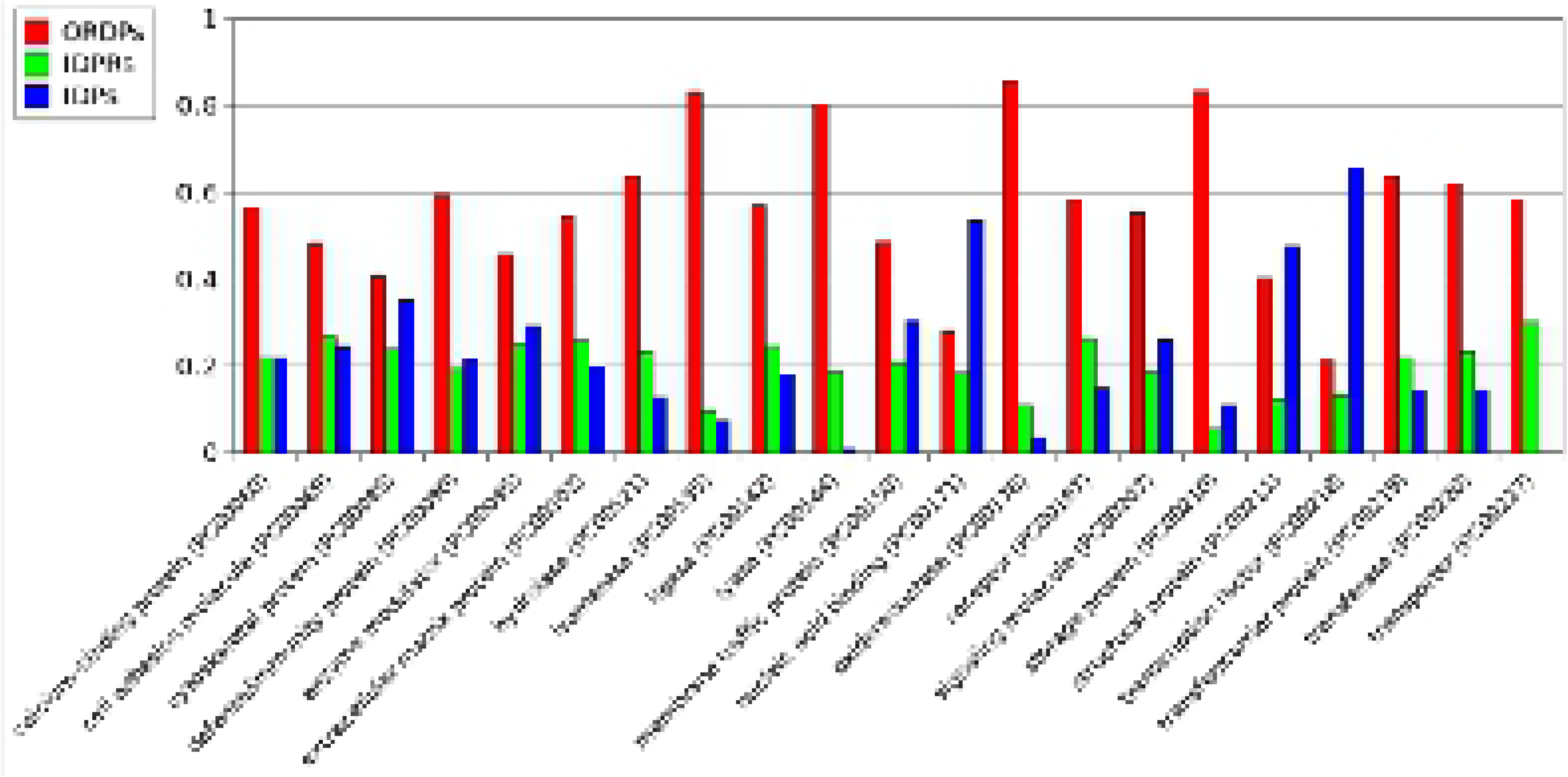
Enrichment of protein functional classes in the protein variants. A functional class is enriched (depleted) in the protein variant that has significantly the highest (lowest) frequency. All the classes shown in this figure passed the enrichment (depletion) test for at least one variant (detailed numerical values can be found in Table in S6 Table).

The results on the enrichment of biological processes, cellular components, and molecular functions in the protein variants are reported in Tables in S7, S8, and S9 Tables, respectively. Many development processes (*pattern specification, ectoderm, mesoderm, endoderm*, and *system development*), chromatin and nucleic acid binding are enriched in IDPs whereas ORDPs enrich *anatomical structure morphogenesis, death*, and *cell differentiation.* Notably, IDPRs enrich only *calcium-dependent phospholipid binding* and *helicase activity*. Finally, IDPs and IDPRs do not enrich any cellular component.

### Over-representation and enrichment of protein variants in diseases

In Fig 4 and Table in S10 Table, we report the over-representation of the protein variants (ORDPs, IDPRs, and IDPs) among the proteins related to cardiovascular diseases, neurodegenerative diseases, and cancer. IDPs are over-represented in cancer and under-represented in cardiovascular diseases. On the contrary, ORDPs are over-represented in cardiovascular diseases and under-represented in cancer. IDPRs are over-represented in all the diseases here considered, and particularly in neurodegenerative diseases.

**Fig 4.**
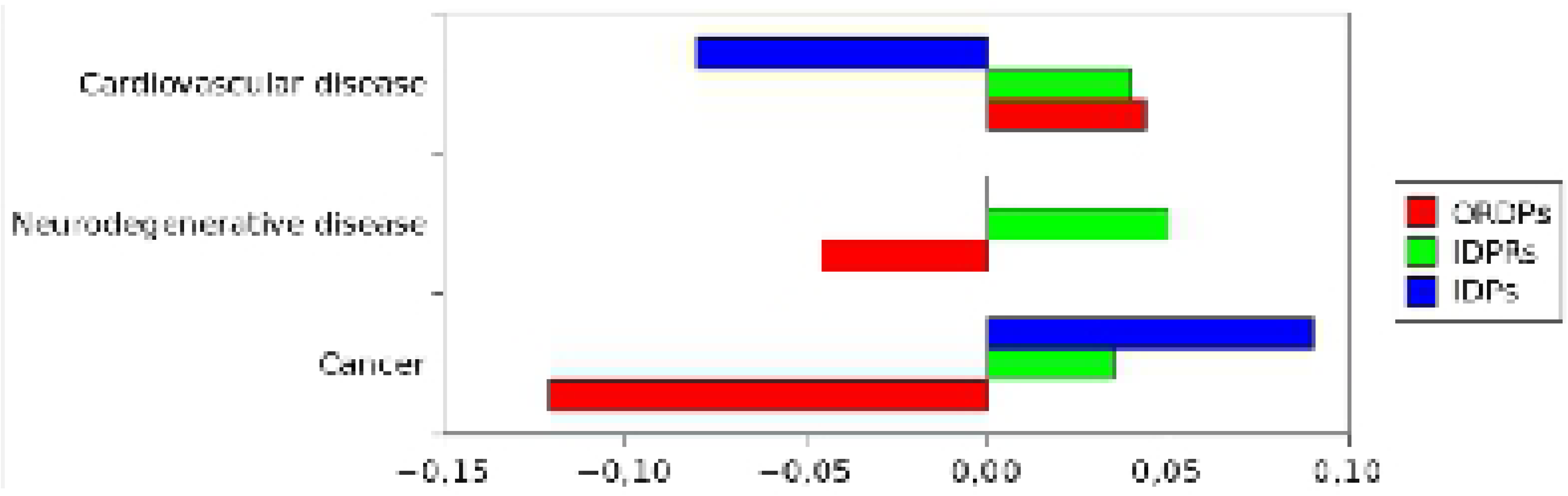
Over-representation of the variants of disorder in diseases. Bar charts of the normalized differential occurrence (with respect to the human proteome) of ORDPs, IDPRs, and IDPs in three groups of disease-related proteins.

In Fig 5 and Table in S11 Table, we report the enrichment of protein variants in diseases. Notably, cancer-related proteins are specifically enriched in IDPs. Both, proteins related to cardiovascular diseases and those related to neurodegenerative diseases are enriched in ORDPs. IDPRs are depleted in all three categories of disease-related proteins (not significantly among proteins related to cardiovascular diseases, see Table in S11 Table).

**Fig 5.**
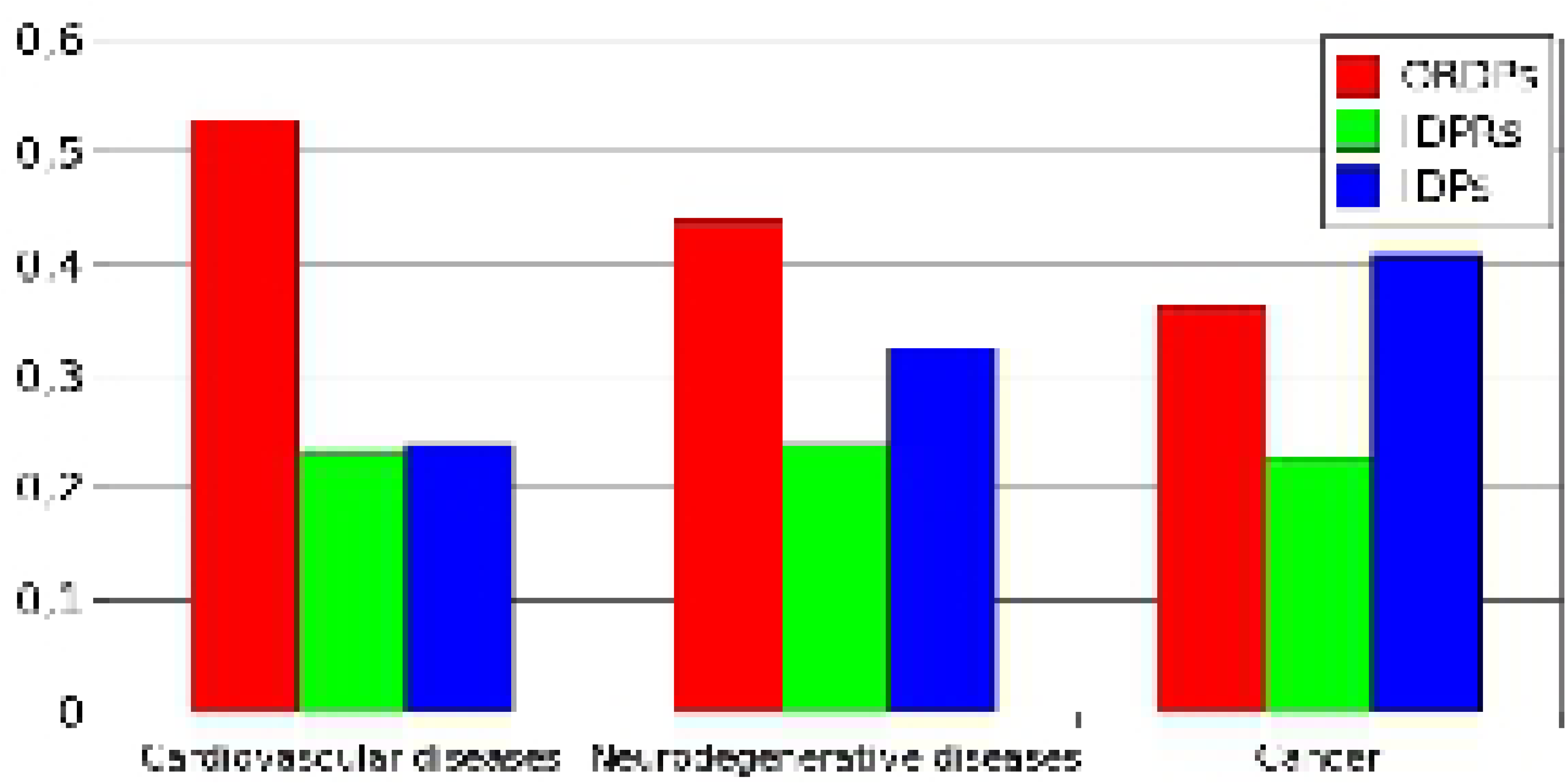
Enrichment of the protein variants in diseases. Different groups of disease-related proteins are enriched (depleted) in the protein variant that has significantly the highest (lowest) frequency. Each group is significantly enriched (depleted) in at least one of the variants (detailed numerical values can be found in Table in S11 Table).

## DISCUSSION

In this study we operationally distinguish the functional spectra and biological roles of three broad variants of intrinsic protein disorder: i) *ordered proteins* (ORDPs), ii) *structured proteins with intrinsically disordered protein regions* (IDPRs), and iii) *intrinsically disordered proteins* (IDPs). The focus of this work is on the functional distinction and the different roles in diseases of IDPs and IDPRs, as we distinguish them.

Many studies aiming at a general classification of functional roles of protein intrinsic disorder were based on the required presence of long disordered regions in the sequence [18-20, 42-45], thus considering IDPRs and IDPs as a single entity [see, for example, 18-20, 42-45].

However, the presence of just one long disordered segment (longer than 30 disordered residues) does not imply that a protein is globally intrinsically disordered, i.e., that it lacks a stable three-dimensional structure. With this criterion, it is impossible to discriminate between IDPs and IDPRs. Moreover, the general requirement of 30 consecutive disordered residues as a fingerprint of IDPs is neither experimentally justified nor universally accepted in the literature. Indeed, this length ranges from 20 residues in Necci et al. [63] up to 50 residues, as in Dunker et al. [2] and Ward et al. [44]. Our distinction is based precisely on this point: the difference between IDPs and IDPRs depends on the relative proportion of the unstructured regions. It has been shown that proteins with more than 30% of disordered residues are more prone to proteolytic degradation [46], likely due to the absence of a well-defined tertiary structure. In line with this argument and in agreement with other studies that adopt the same threshold [61,64], we have then defined proteins with more than 30% of disordered residues as IDPs. Finally, to identify IDPRs, we considered the study by van Der Lee et al. [14]. They report that a protein has a shorter half-life (due to the protease-vulnerability of its disordered regions) either if it has a segment of 30 consecutive disordered residues located at the C- or N-terminus or a segment of 40 consecutive disordered residues in a generic position of the polypeptide chain. Moreover, unstructured regions longer than ∼25 residues have been considered to evolve, function, and exist as independent, disordered domains [15,65,66]. Thus, we identified IDPRs as those proteins with less than 30% of disordered residues but with at least one disordered domain.

The main result of this study is that intrinsically disordered proteins (IDPs) and structured proteins with intrinsically disordered protein regions (IDPRs) have different functional roles in the cell. We found that IDPs are mainly associated with nucleic acid binding proteins, chromatin binding proteins, transcription factors, and many developmental processes. The functional classes in which IDPRs are over-represented substantially overlap with those in which ORDPs are also over-represented (transporters, receptors, and catalytic activities). This outcome shows that the functional roles of IDPRs are more similar to those of ORDPs than to those of IDPs. Moreover, IDPs are specifically over-represented in PANTHER functional classes and GO annotations where both IDPRs and ORDPs are under-represented. We conclude that the broadly shared statement that IDPs are specifically involved in signaling and cell-regulation [67] should be taken with caution. As regards to disease-related proteins, we observe that IDPs are abundant among cancer-related proteins but not among proteins associated with cardiovascular and neurodegenerative diseases, where ORDPs seem to dominate. This observation better specifies the quite established D2 concept [32].

Overall, our approach is useful to discriminate between IDPs and IDPRs operationally, and we have used it to demonstrate the different functional roles of these two variants in the human proteome. As a conclusion, we believe that it is important to correctly evaluate the occurrence of disorder in proteomes, in evolutionary studies, in understanding functional aspects related to different flavors of protein disorder, and in deciphering the role of these variants in the emergence of diseases.

## ABBREVIATIONS

ORDPs: ordered proteins;
IDPRs: structured proteins with intrinsically disordered regions;
IDPs: intrinsically disordered proteins.

